# STRkit: precise, read-level genotyping of short tandem repeats using long reads and single-nucleotide variation

**DOI:** 10.1101/2025.03.25.645269

**Authors:** David R Lougheed, Tomi Pastinen, Guillaume Bourque

**Author notes:** Corresponding authors: David R Lougheed and Guillaume Bourque.

## Abstract

Variation in short tandem repeats (STRs) is implicated in Mendelian disease and complex traits, but can be difficult to resolve with short-read genome sequencing. We present `STRkit`, a software package for genotyping STRs using long read sequencing (LRS) that uses nearby single-nucleotide variants to improve genotyping accuracy without a priori haplotype information. We show that `STRkit` has unique strengths versus other methods: it can use data from both major LRS technologies (Pacific Biosciences HiFi [PB] and Oxford Nanopore [ONT]) to output both allele and read-level copy number and sequence, performs best in benchmarking with F1 scores of 0.9633 and 0.9056 with PB and ONT data respectively, achieves a Mendelian inheritance rate of 97.86% with PB data, and is open source software. `STRkit`’s features open up new possibilities for association testing, assessing patterns of STR inheritance, and better understanding the functional effects of these notable repeat elements.

## BACKGROUND

Short tandem repeats (STRs), or microsatellites, are repetitive genomic elements found in both eukaryotes and prokaryotes where a small motif, from 1 or 2 to 6-13 base pairs long (Lander *et al*. 2001, Ellegren 2004, Chiu *et al*. 2021), is repeated several times in a row. These regions cover around 3% of the human genome (Lander *et al*. 2001), are often multi-allelic, and are significant contributors to total genomic variation (Ellegren 2004, English *et al*. 2024, Liao *et al*. 2023, Hannan *et al*. 2018). STRs typically have high rates of mutation, usually via stepwise repeat expansion or contraction (Fan & Chu 2007, Ellegren 2004).

Variation in STRs is associated with over 60 inherited and sporadic diseases in humans (Gall-Duncan *et al*. 2022). In Huntington’s disease, fragile X syndrome (FXS), and others, copy number expansion beyond a threshold causes the disease, and further expansion is associated with phenotypic severity (Igarashi *et al*. 1992; Duyao *et al*. 1993; Brinkman *et al*., 1997; Matsuura *et al*., 2000; Blauw *et al*. 2012; Bragg *et al*. 2017). The relationship between copy number and disease severity is not always linear; for example, with fragile X-associated primary ovarian insufficiency, peak severity occurs at an intermediate copy number (Allen *et al*. 2021). STR motif composition can also affect phenotype: insertion of a stretch of a non-canonical motif into a non-coding STR in the DAB1 gene is responsible for a form of spinocerebellar ataxia (Seixas *et al*. 2017). Beyond expansion detection, precisely determining copy number and motif composition in disease-causing STR expansions may be required to fully quantify their relationship with phenotype.

Other phenotypic relationships with STRs are not limited to a single locus: autism is associated with a genome-wide increase in rare STR expansions (Trost *et al*. 2020) and small *de novo* STR mutations (Mitra *et al*. 2021). Expansion and somatic instability in STRs feature in several cancers (Erwin *et al*. 2022, Yamamoto & Imai 2019). STRs are implicated in gene expression and regulation (Gymrek *et al*. 2016; Hannan, 2018; Cui *et al*. 2025), including by acting as binding sites for transcription factors (Horton *et al*. 2023). Direct STR genotyping at a genome scale is thus valuable; STRs cannot always be imputed using SNVs (Gymrek *et al*. 2016, Saini *et al*. 2018), as SNV-STR linkage disequilibrium (LD) is lower than SNV-SNV pair LD (Willems *et al*. 2014, Press *et al*. 2018). Examination of STR variation in the human genome may help to address the “missing heritability” problem, explaining a portion of complex trait heritability not yet identified (Hannan 2018).

However, STRs are difficult to characterize with short read sequencing (SRS) technologies due to often long repeat tracts and lack of flanking sequence within reads (Narzisi & Schatz 2015, Mousavi *et al*. 2019), affecting mappability to a reference genome. Some SRS STR genotyping tools can estimate larger-than-read-length allele sizes (Dolzhenko *et al*. 2017, 2019, Mousavi *et al*. 2019) but cannot capture variation in motif structure nor precisely genotype these long expansions (Mousavi *et al*. 2019). By contrast, modern long read sequencing (LRS) technologies like Oxford Nanopore’s nanopore sequencing (ONT) with newer chemistry, or Pacific Biosciences (PacBio) high-fidelity circular consensus sequencing (HiFi), can give accurate reads of sizes in the tens of kilobases (Gustafson *et al*. 2024; Tanudisastro *et al*. 2024; Wenger *et al*. 2019). These reads span pathogenic expansions that exceed SRS read length (Hannan 2018) and include more flanking sequence. These flanking sequences may include single-nucleotide variants (SNVs) that could be used to cluster STR-spanning reads into alleles, as SNVs are the most reliably-called form of variation with LRS (Wenger *et al*. 2019, Gustafson *et al*. 2024).

Several LRS STR genotyping tools have been recently developed, including `Straglr` (Chiu *et al*. 2021), `TRGT` (Dolzhenko *et al*. 2024), `LongTR` (Ziaei Jam *et al*. 2024), and `STRdust` (De Coster *et al*., 2024). These methods take standard BAM/CRAM alignments as input and output variants as VCF files. `LongTR` and `TRGT` have been tested on whole-genome catalogs of STRs, meaning they should be suitable for association testing and novel expansion discovery, among other whole-genome-scale applications. However, they do not take full advantage of LRS data, or have additional restrictions which limit their use – `TRGT` can use nearby SNVs to aid genotyping, but is license-restricted to only work with PacBio data; `Straglr` does not output consensus sequences for alleles; and neither `LongTR` nor `STRdust` output read-level data, limiting their use for motif composition analysis or detecting within-allele somatic instability.

Here, we introduce a new STR genotyping software, `STRkit`, designed to operate primarily with accurate LRS data. `STRkit` can optionally call proximate SNVs and use them to cluster and locally phase STR alleles without needing a phased SNV call-set a priori. Data can be output at the read level, enabling analysis of intra-allele STR copy number and/or motif composition to analyse somatic instability. We also evaluate `STRkit`’s performance for STR genotyping, and demonstrate the value of incorporating SNVs into the process by benchmarking `STRkit` against existing LRS STR genotyping methods using the Genome-in-a-Bottle (GIAB) tandem repeats benchmark (English *et al*. 2024) for the HG002 individual of the well-characterized GIAB Ashkenazi trio. We evaluated performance on an STR-only subset of this benchmark using `Truvari` (English *et al*. 2022) and `Laytr` (English *et al*. 2024), and by assessing rates of Mendelian inheritance in trio genotypes. Finally, we demonstrate how read-level STR genotype information produced by our tool replicates a known pattern of somatic repeat instability in targeted sequencing of a pathogenic expansion.

## RESULTS

### The STRkit software package

We created an STR genotyping method, `STRkit`, as an easily-installable Python package. Our algorithm takes aligned long-read sequencing (LRS) BAM or CRAM files as input, with an optional realignment step for mis-mapped reads (**Figure 1A** and **Methods**). It then determines STR allele copy number and optionally genotypes nearby SNVs, whose locations are extracted from a user-provided VCF file. Depending on what SNV data are present, `STRkit` either uses a Gaussian mixture model allele-calling approach with copy numbers, segregates alleles based on SNV haplotype groups, or uses a mix of both information sources. If two STRs have common SNV haplotype groups, they will be locally phased. If a phased alignment file is provided, `STRkit` can use read haplotype data instead. Genotypes are then reported in either VCF, TSV, or JSON formats. JSON output can be explored in an included web-based visualization tool (**Figure 1B**). Our software package also includes a Mendelian inheritance (MI) analysis tool, for benchmarking callers or detecting candidate *de novo* mutations. We compared the features of `STRkit` against those of other LRS STR genotyping packages to highlight its unique features, including the MI tool, SNV incorporation and output during STR genotyping, and read-level copy number output (**Table 1**).

**Figure 1:**
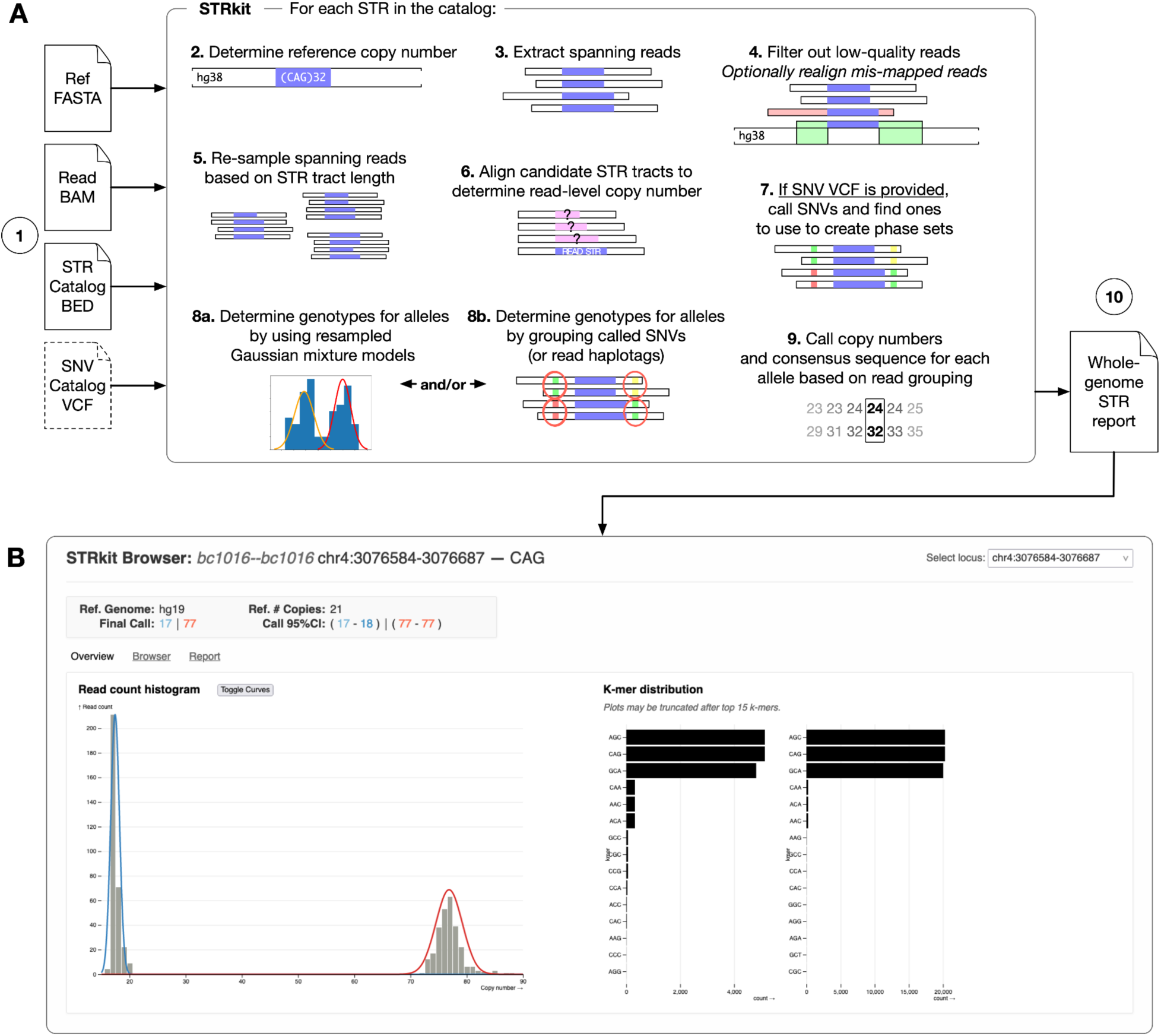
Flowchart of the `STRkit` genotyping and visualization workflow. **A.** The genotyping workflow (also see **Methods**). Our approach includes an optional SNV incorporation step, which can find heterozygous SNVs proximate to STRs and use them to cluster reads to call STR alleles. **B.** Users can explore the generated report in the `STRkit` visualization tool. The read count histogram shows the read-level distribution of copy numbers for the locus. Here, a pathogenic expansion in the HTT gene is shown. The k-mer distribution plots show motif-sized k-mer sequence diversity among read STR sequences for each allele peak.

**Table 1:** Feature comparison of long-read sequencing STR genotyping tools.

| Feature | STRkit | LongTR | TRGT | Straglr | STRdust | Notes |
| --- | --- | --- | --- | --- | --- | --- |
| Copy number output | ✓ | - | ✓ | ✓ | - |  |
| Allele size confidence intervals | ✓ | - | ✓ | ✓ | - |  |
| Allele consensus sequence output | ✓ | ✓ | ✓ | - | ✓ |  |
| De novo proximate SNV phasing | ✓ + output | - | ✓ | - | - | STRkit and TRGT can use heterozygous SNVs to cluster STR alleles; STRkit can output them to VCF. |
| Existing phased SNV incorporation | - | ✓✓ | - | - | - |  |
| Haplotagged alignment file support | ✓ | ✓ | ✓ | - | ✓ | All but Straglr can use phased read data from, e.g., WhatsHap (Martin *et al.* 2016) to call STRs. |
| Methylation handling | - | - | ✓✓ | - | - |  |
| ONT read support | ✓ | ✓ | - | ✓ | ✓ | TRGT: Theoretically works on ONT data, but forbidden by software license. |
| Read-level data | ✓ | - | Partial | ✓ | ✓ | STRkit: JSON output with read-level peak ID + sequence data; TRGT: overlapping reads in BAM; Straglr: TSV output with read-level copy numbers; STRdust: read-level sequence output in VCF. |
| Mendelian inheritance calculation tool | ✓✓ | - | - | - | - | STRkit includes a tool to output loci which do not respect Mendelian inheritance in a set of trio VCFs. |
| Free and open-source software license | Yes (GPLv3) | Yes (GPLv2) | - | Yes (GPLv3) | Yes (MIT) | TRGT's license restricts it to be only used with PacBio sequencing data, and the software cannot be forked and subsequently re-distributed. |
| Multi-threading/processing | ✓ | - | ✓ | ✓ | ✓ |  |
| Pre-built Docker image | ✓✓ | - | - | - | - | STRkit is available as a pre-built Docker image, which can be incorporated into, e.g., Nextflow/WDL pipelines. |

Comparison of features of STRkit with those of recent long-read sequencing STR genotyping tools: LongTR, TRGT, Straglr and STRdust. Features with double check marks are unique to that particular tool.

### Precise STR genotyping with long reads using proximate SNVs

To evaluate the performance of `STRkit`, we first obtained PacBio HiFi reads at ∼32x coverage and ONT R10 Duplex reads at ∼12x coverage for the Genome-in-a-Bottle (GIAB) HG002 individual from the GIAB Ashkenazi trio, and aligned them to the HG38 reference genome (see **Methods**). We then genotyped 914 676 short tandem repeat regions derived from a ∼1.7 million tandem repeat benchmark variant set developed for this individual (English *et al*. 2024) and released as an official benchmark by the GIAB consortium. The benchmark covers ∼8.1% of the HG38 human reference genome, but not all of these benchmark regions are STRs; some have homopolymers or larger TRs, which we excluded to provide more precise estimates of performance (see **Methods**). We also removed regions with more than one tandem repeat because the tool we used to evaluate performance on the benchmark, `Truvari` (English *et al*. 2022), cannot handle proximate unphased variants. Our final set of benchmark STRs covers ∼0.87% of HG38. Separately, we confirmed the accuracy of `STRkit`’s local SNV phasing step by comparing it to alignment files for the Ashkenazi trio, which were haplotype-tagged using `DeepVariant` (Poplin *et al*. 2018) and `WhatsHap` (Martin *et al*. 2016). Across the trio, we found a phase error rate of approximately 0.12% in phase sets with more than one locus.

To evaluate `STRkit` against other tools, we also genotyped our benchmark set of STR loci using `LongTR` (Ziaei Jam *et al*. 2024), `Straglr` (Chiu *et al*. 2021), `STRdust` (De Coster *et al*. 2024), and `TRGT` (Dolzhenko *et al*. 2024). We then assessed their respective performance using `Truvari` and `Laytr` (English *et al*. 2024); except for `Straglr`, whose output is not compatible with `Truvari`. These results are summarized in **Table 2**. With HiFi data, `STRkit` performed best overall in most metrics (accuracy = 0.9767, F1 = 0.9633), although `TRGT` achieved marginally better recall (0.9697 versus 0.9552 for `STRkit`). With ONT data, `STRkit` (accuracy = 0.9438, F1 = 0.9056) performed better than `LongTR` (accuracy = 0.9296, F1 = 0.8786) and `STRdust` (accuracy = 0.8443, F1 = 0.7693) in all metrics.

**Table 2:**
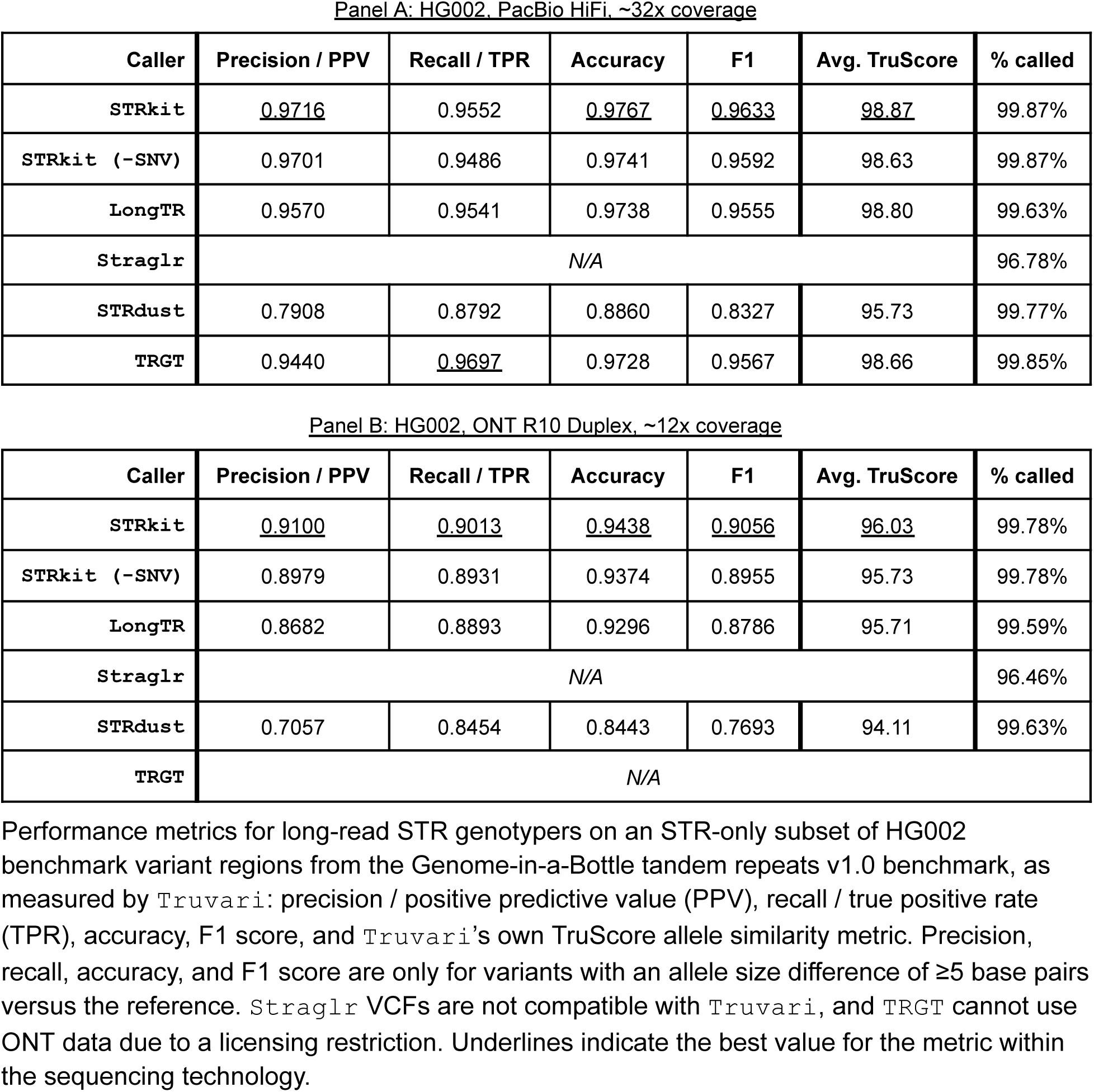
Performance metrics for long-read STR genotypers on the HG002 benchmark subset <u>Panel A: HG002, PacBio HiFi, ∼32x coverage</u>

`Truvari` also reports a compound variant quality score, called the “TruScore”, combining overlap, size similarity, and sequence similarity between a given call and the benchmark into an overall score (English *et al*. 2022). With HiFi data, `STRkit` achieved the best average TruScore of 98.87, closely followed by `LongTR` with 98.80 and `TRGT` with 98.66 and beating `STRdust`’s average score (95.73) by a larger margin. With ONT data, a similarly small average TruScore difference was observed between `STRkit` (96.03) and `LongTR` (95.71), followed again by `STRdust` (94.11).

Using the `Laytr` tool, we stratified performance metrics (F1 score, precision, recall) by the locus-maximum absolute change (Δ) in allele size from the reference genome (**Figure S1**). Most STR Δ allele sizes are small (95.5% of allele size changes are within 50 base pairs (bp) of the HG38 reference genome) and that is where `STRkit` with HiFi data shows a slight but measurable improvement over other tools in terms of F1 score and precision. With ONT data for small and intermediate allele size changes versus the reference (within 200 bp of HG38), `STRkit` performed markedly better versus other tools. With larger Δ allele size, all tools performed worse with both sequencing technologies, with `TRGT` appearing to perform the best with these loci when it could be used (PacBio only). It was with these larger Δ allele sizes where `STRkit`’s SNV incorporation helps its genotyping performance the most.

Our evaluation of `STRkit` without the SNV incorporation step confirmed that using SNV data improves STR genotyping. All metrics output by `Truvari` for `STRkit` improved when the SNV step was included, for both HiFi and ONT data (**Table 2**). For most loci in the call-set (∼72% across the trio) `STRkit` used information from at least one heterozygous SNV for clustering reads into alleles (**Table S1**).

### High-quality genotypes track STR allele inheritance in trios

`STRkit` also includes a tool for calculating the rate of Mendelian inheritance (MI) of alleles from parents to offspring, in terms of copy number and allele sequence/length. Loci are considered to follow MI if allele sequences from the child can be found in the parents. This process also identifies loci that do not meet expectations of MI in the trio – sites of potential *de novo* mutation. Versions of this metric have been used to evaluate other STR genotyping software, e.g., `GangSTR` (Mousavi *et al*. 2019). On the Ashkenazi trio, `STRkit` achieves a sequence-wise (seq.) MI rate of 97.85% (98.07% in terms of seq. length, 99.08% in terms of seq. length ±1 base pair). Without SNVs, `STRkit` achieves a lower seq. MI rate of 93.65% (95.68% seq. len., 98.43% ±1bp), indicating that incorporating SNVs to cluster reads improves allele sequence quality. `LongTR` and `TRGT` again perform almost as well; `LongTR` achieves a seq. MI rate of 94.52% (94.85% seq. len., 97.54% ±1bp), and `TRGT` achieves a seq. MI rate of 94.68% (97.10% seq. len., 98.98% ±1bp). `Straglr` and `STRdust` achieve significantly lower rates of MI. These results are summarized in **Table S2**.

We then stratified sequence-wise MI by reference genome locus length (**Figure 2**). The bulk of loci are small, which is again where `STRkit` has the clearest advantage over other methods when incorporating SNVs. Above ∼200 base pairs in length, the rates drop off, as STRs become harder to sequence accurately and are more likely to accumulate mutations in their passage from parent to child.

**Figure 2:**
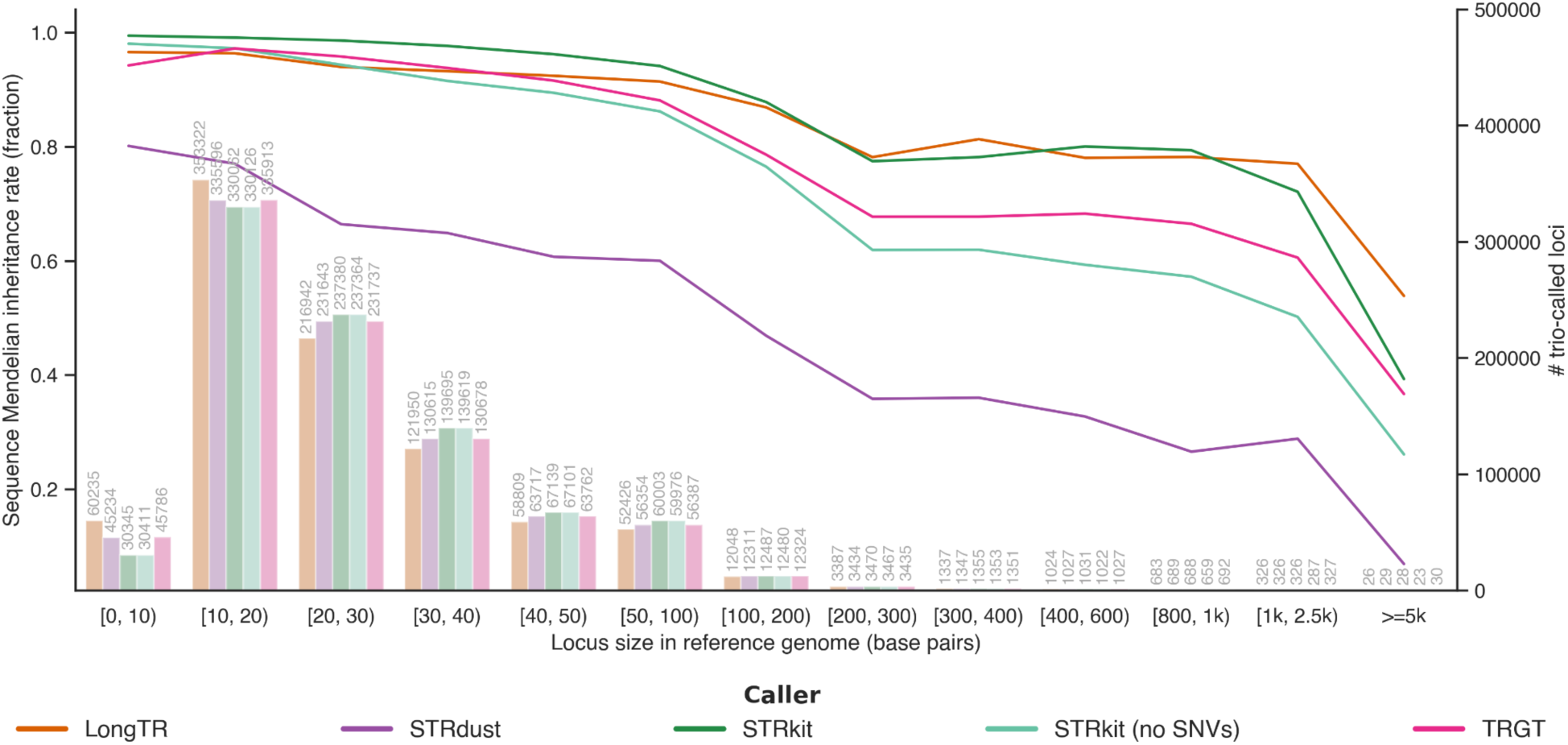
Rates of sequence Mendelian inheritance (MI), and number of trio-called loci, by reference genome locus length. Expectations of MI are met if both child allele sequences can be seen in the two parents (one in the mother and one in the father). The vast majority of short tandem repeat regions in the catalogue are below ∼200 base pairs long, where `STRkit` (with SNV incorporation) achieves the highest rate.

### Computational performance of SNV-informed STR genotyping

Most genotyping tools tested (`LongTR`, `STRkit`, and `TRGT`) can analyze the filtered HG002 benchmark catalogue in under a day measured in core-minutes using ∼32x coverage PacBio HiFi data and ∼12x ONT R10 Duplex data (**Table S3**). `STRkit` is slower than `TRGT` and `LongTR`, but is on the same order of magnitude (in the 100s of core-minutes); in contrast, `Straglr` is in the 1000s of core-minutes, and `STRdust` is the slowest at ∼10 000 core-minutes on average for the GIAB trio individuals sequenced with HiFi. Where possible, we used 8 cores of an Intel Gold 6148 processor; `LongTR` does not support using multiple CPU cores, but is the fastest in terms of core-minutes used (**Table S3**). `STRkit` was the fastest multi-core program for ONT. `STRkit` consumes more memory than `TRGT` or `LongTR` with HiFi data on average, but less than `STRdust` and `Straglr`, and the SNV incorporation step increases its memory usage. `STRkit` without SNV incorporation is the most memory-efficient STR caller for ONT.

### Read-level genotyping of targeted long-read sequencing data captures known instability in a pathogenic expansion

We ran `STRkit` and other tandem repeat callers on targeted CCS sequencing data of pathogenic repeat expansions in the HTT and FMR1 genes (**Table S4** and **Methods**). We ran `STRkit` using its targeted sequencing mode, without SNV incorporation as the targeted sequencing data lacks sufficient flanking sequence. The callers successfully detected the repeat expansions in almost all cases, except for one HTT expansion missed by `Straglr`, and one FMR1 expansion missed by `STRdust`. The genotypes overall were highly concordant.

`STRkit`, `Straglr`, and `TRGT` output copy numbers directly, and the former two have read-level copy number output, allowing further expansion analysis. De Luca *et al*. (2021) found that one of these samples, NA20253, has three copy number peaks, i.e., mosaic alleles, which they measured at 107, 134, and 175 CAG repeats, with the first being the most prominent. **Figure 3** shows the the read-level data for this expanded allele from `STRkit`’s output, along with annotations showing the peaks found by De Luca *et al*. (2021); we observe three read peaks which mirror those found using their triplet repeat-primed PCR assay, but slightly shifted towards a higher copy number.

**Figure 3:**
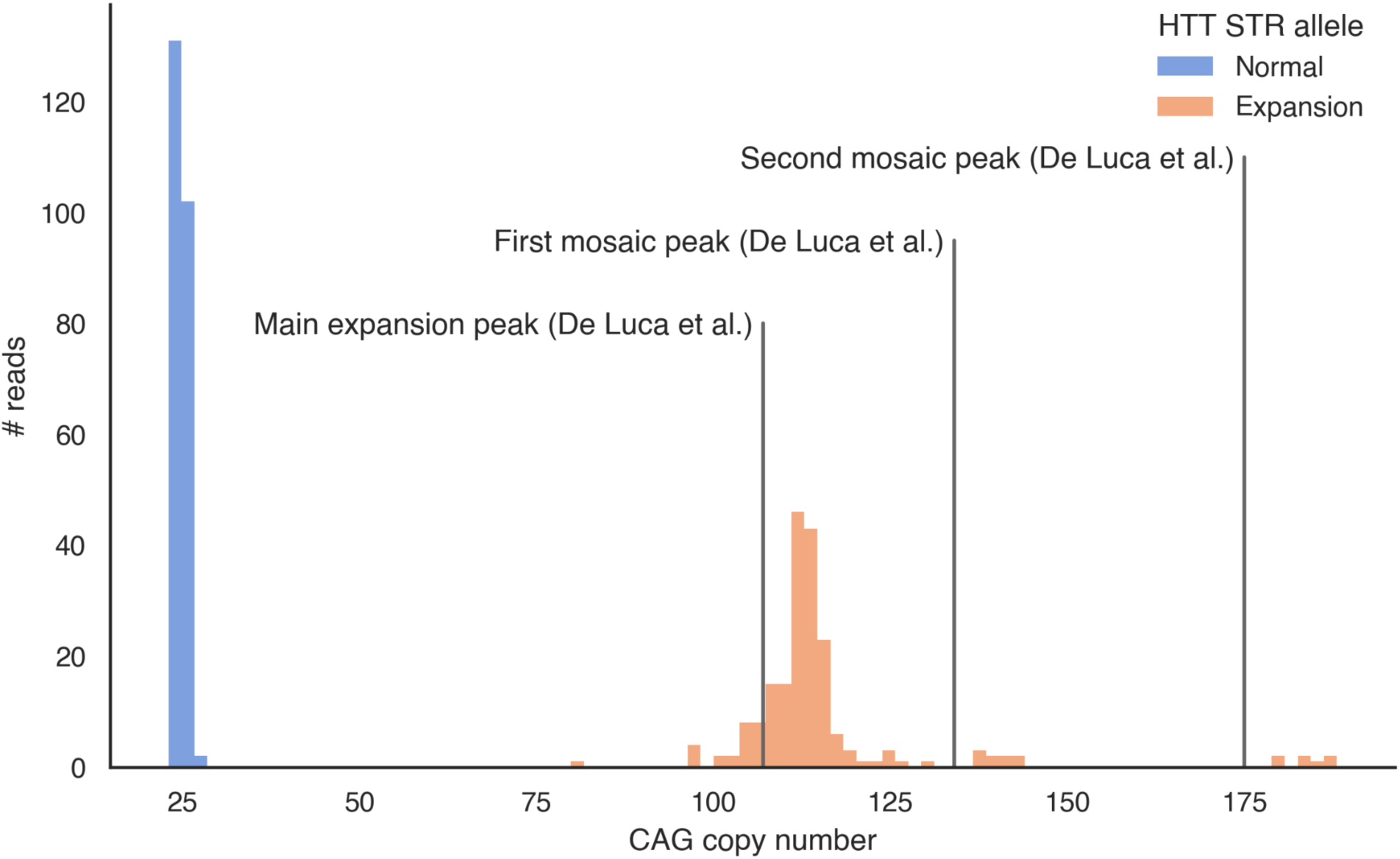
HTT allele copy number distribution in targeted sequencing of the NA20253 sample. Vertical lines show the three expansion copy number peaks found by De Luca *et al*. (2021), which fall near read peaks in `STRkit` output – two mosaic alleles, at 134 and 175 repeats, with the main allele peak (with the majority of reads) at 107 repeats.

## DISCUSSION

Here, we present `STRkit`, a new tool for calling short tandem repeat genotypes from long read sequencing (LRS) data. Our tool has a unique combination of features, filling a niche in the STR genotyping analytical space: an STR caller which can operate at a genome-wide scale with both major LRS technologies, precisely genotype both sequence and copy number, incorporate proximate SNVs into the calling process and use them to phase STRs locally, and report read-level data (**Table 1**). Both STR copy number and sequence are phenotypically relevant properties (Olson *et al*. 2023); however `LongTR` and `STRdust` only output sequence, and `Straglr` outputs only copy number. Of the tested tools, all are LRS technology-independent except `TRGT`, which can only be used with PacBio data due to a licensing restriction.

Technology-independent tools are critical, because they enable repeatable research and community-inclusive development, allow for confirmation of findings across technologies, and allow the best genotyping approach to be evaluated and applied regardless of data source – for example the cross-technology benchmarking we performed to evaluate STRkit.

Our benchmarking with `STRkit`, when incorporating proximate SNVs into the genotyping process, demonstrates state-of-the-art performance in several metrics with both PacBio HiFi data and modern nanopore sequencing data (ONT R10 Duplex). While performance gains are modest with HiFi data versus the next best-performing tool, `LongTR` (**Table 2**), maximizing precision and accuracy has direct benefits for genome-wide analysis, where statistical power in correlation tests would be reduced by lower-quality genotypes. With ONT data, our performance gains were larger, achieving a ∼6% higher F1 score than `LongTR` and narrowing the performance gap between 30x HiFi and 12x ONT data. Our SNV incorporation process allows clustering STR-overlapping reads into alleles without relying solely on copy number. This aids our genotyping approach to a small but measurable degree (**Table 2**), and for most loci we can incorporate SNV information into the STR calling process (**Table S1**), although it may not be the primary explanation for `STRkit`’s improved benchmark metrics versus other tools, especially with ONT data (**Table 2**, **Figure S1**). SNV data captured by `STRkit` may prove useful for more than genotyping; in downstream analysis, they could aid, for example, trio phasing or STR mutation modeling, although our current approach only calls heterozygous STR-proximate SNVs, and we have not extensively validated these calls against dedicated SNV callers like `bcftools` (Danecek *et al*. 2021).

When metrics are faceted by allele size difference (Δ allele size) versus the reference genome, tools generally performed well until Δ allele size reached ∼2.5k base pairs or longer (**Figure S1**), except for a noticeable dip in genotyping quality in the the 3-500 base pair range. Manual inspection indicates that some of these may be regions with non-canonical motifs or non-tandem-repeat sequences embedded within a tandem repeat locus, which may impact `STRkit`’s modeling of candidate repeat tract sequences or result in reads being filtered out to a greater degree versus other tools.

Of the five callers, `STRkit`, `LongTR`, and `TRGT` best accommodated the large catalogue of ∼900 000 loci, with relatively quick runtimes (in the 100s of core-minutes) and low memory requirements (**Table S3**). Our general approach to genotyping does add a computational overhead, as it is slower to genotype the benchmark catalogue versus `LongTR` and `TRGT`. We believe that this is due to our additional read-level data collection and copy-number determination approach. The SNV incorporation step in `STRkit` appears to speed up genotyping (654 versus 978 core-minutes on average in the trio), indicating that future performance improvements could come primarily from optimizing other procedures (e.g., read processing or copy number determination) within our method. `STRdust` and `Straglr` both had significantly longer runtimes (1000s of core-minutes), and `Straglr` used an order of magnitude more memory than the other tools. `STRkit` is the only one of the three most performant tools (in terms of both genotype quality and computation speed) which can report full read-level copy number data, meaning it may be useful for genome-wide somatic instability detection. `LongTR`’s lack of parallel processing support means it avoids parallel processing overhead, at the cost of longer per-sample runtime, whereas other methods were evaluated using eight CPU cores, requiring inter-process communication. `STRkit`’s shared SNV context, used for local phasing of STR genotypes, likely results in slower-than-linear performance gains with more CPU cores, trading off additional computation for increased genotype quality.

In our trio analysis for Mendelian inheritance (MI), we quantified the proportion of STR loci where both alleles in the HG002 child were found in the parents. `STRkit` had the best performance across the three versions of the metric we tested (**Table S2**). MI is limited as a benchmarking metric, however, because systematic errors such as off-by-one errors which affect both parent and child may not be adequately considered. Despite this, our high MI rates, when combined with high absolute scores on the benchmark, indicate that `STRkit` should be the most accurate currently-available tool for genome-scale STRs genotyping, and potentially for detecting the most common category of *de novo* STR mutation events: ±1 repeat unit changes. However, without a more complete mutation detection procedure with statistical confidence measures to control for false discovery, we cannot claim that all loci which do not respect MI are true *de novo* mutations.

So far, we have discussed genome-wide benchmarking results across ∼900 000 tandem repeat loci. Very few STR loci have known occurrences of pathogenic expansions – Gall-Duncan *et al*. (2022) reported 63 as of 2021. `STRkit` provides a unique feature-set useful for close examination of specific expansion loci. To demonstrate this, we used its targeted sequencing mode to genotype STRs with known expansions in the HTT or FMR1 genes (**Table S4**), where, like other tools, it correctly detects the expanded allele. Unlike other tools, `STRkit` can both generate a whole-allele consensus sequence and report read-level copy number and sequence data, including motif-sized k-mer distributions, at the same time (**Figure 3**). `STRkit` does not directly detect somatic instability, unlike a tool such as `prancSTR` (Sehgal *et al*. 2024); however, `STRkit`’s read-level copy number data replicates a previously-captured multi-peak HTT mosaic expansion in a cell line sample (De Luca *et al*. 2021). Our peaks are slightly offset from the reported peaks in this expansion, possibly due to the passage of the sampled cell line. This pattern revealed itself only with high-depth targeted sequencing: this may be critical for fully characterizing expansion alleles.

Beyond somatic instability detection, there are other STR genotyping features `STRkit` does not yet implement. Methylation reporting using HiFi’s base methylation data, for example, remains exclusive to `TRGT`. Our calling approach also has limitations: reads are primarily clustered based on motif copy number if SNV or haplotype information is not available; this is useful for obtaining copy number confidence intervals, but may miss more subtle allelic sequence variation. In the absence of SNV data, our tool may also miss variation when faced with large allele differences versus the HG38 reference genome, to a greater degree than other methods (**Figure S1**). Finally, `STRkit`, like most of the other tools tested, is a catalogue-based genotyper – it requires a list of loci with reference genome coordinates and a motif. This limits what can be genotyped to only what is already present in the reference, and may not function well with non-reference motifs within known loci or may miss certain disease-causing expansions completely (Tanudisastro *et al*. 2024). Annotating an assembly or using a *de novo* approach such as `ExpansionHunter Denovo` (Dolzhenko *et al*. 2020) or `Straglr` without a provided catalogue can find repeats that `STRkit` and others cannot.

## CONCLUSIONS

Short tandem repeats (STRs) cover around 3% of the human genome, are implicated in genetic disorders and transcriptional regulation, and are major contributors to overall genomic variation. STRs are important to genotype directly, as they cannot always be imputed with single-nucleotide variation (SNVs), but these elements are difficult to resolve with traditional sequencing methods and complex to analyze – they are multi-allelic and defined by both their length and motif composition, rather than a simple binary genotype. `STRkit` uses accurate long-read data and a strategy which incorporates both SNVs and copy-number genotyping into the STR genotyping process to achieve better performance in several metrics. The tool also includes read-level copy number and motif composition data for these loci, while many other methods do not. The approach fills a niche in the STR genotyping software ecosystem and opens up new possibilities for association testing, assessing patterns of STR inheritance, and better understanding the functional effects of these repeat elements.

## METHODS

### STR genotyping approach

The general genotyping approach used in STRkit is as follows:

1. The user provides a BAM or CRAM-formatted long-read sequencing (LRS) alignment file, generated using an aligner such as `minimap2` (Li, 2018), as well as a BED-formatted catalogue of STRs to genotype. Optionally, they also provide a VCF with a list of SNVs to genotype if proximate to an STR.
2. For each STR in the catalogue, we optionally re-determine the copy number of the STR in the reference genome to allow for imperfect reference motif copies and adjust the catalogue coordinates to correct for small boundary errors.
3. The set of all reads spanning each STR region (including flanking sequences on either side for realignment) is extracted using the *PySAM* library (v0.22.0; https://github.com/pysam-developers/pysam).
4. Low-quality or badly-aligned reads are filtered out. If realignment is enabled, badly-aligned reads may be realigned to the reference genome.
5. For each read, a series of candidate STR tracts of varying copy number, starting with a candidate closest in length to the aligned read’s STR segment size, are generated using the catalogued motif. Each of these candidate STR tracts are aligned to the read using the *parasail* library’s semi-global alignment algorithm (Daily 2016), incorporating 5’ and 3’ flanking sequences to ‘anchor’ the STR candidate and to allow for complete STR tract deletion. The best-scoring alignment candidate is kept for each read in question. After we have finished this process, we have approximate copy numbers for each read. *If the read-level k-mer functionality is enabled, motif-sized k-mers are computed for each read*.
6. **If an SNV VCF catalogue is provided:** For each read, SNVs that differ from the reference are collected; these are then filtered to those which occur in many reads and can split the reads into two groups, i.e., likely heterozygotic.
7. For each read, a re-sampling weight is generated, corresponding to the estimated inverse probability of observing a read encompassing an STR tract of the same size. This process adds weight to reads with longer STR tracts. Given random read localization on a reference and a read size distribution, longer STR alleles are less likely to be entirely ‘captured’ by a read; in this process, allele size bias is corrected for, and long expansion tracts can be better captured. Reads (with associated copy numbers and SNV calls) are then resampled a number of times (defaulting to 100), according to the re-sampling weight.
8. From here, multiple possibilities are available to call allele peaks:

a. **If there is haplotype information in the alignment file and `--use-hp` is specified:** The haplotype information for the reads is used if available. Otherwise, one of the next three methods is used.
b. **If there is a lot of informational SNVs for the locus:** SNVs are used to split the reads into allele peaks, since SNV assessment is generally very accurate in NGS data. A distance matrix is calculated, and an agglomerative clustering function from the *scikit-learn* library (Pedregosa *et al*., 2011) is applied to group the reads into alleles.
c. **If there is some SNV information available:** SNV data are used, but copy number is incorporated into a compound distance function, where SNV differences are weighed more heavily (5:1 or 10:1) versus copy number differences. Agglomerative clustering is used here as well.
d. **If no or limited SNV information is available:** A Gaussian Mixture Model (GMM) from *scikit-learn* is applied to each re-sampling to derive estimates of allele copy numbers via a bootstrap process. Each allele starts with a two-peak GMM; if one peak is assigned very little explanatory weight after fitting, we switch to a single-peak GMM (i.e., a homozygous model). If the organism is diploid, reads are assigned to peaks based on SNV genotypes (if available) or bootstrapped estimates of peak mean, standard deviation, and weight – the complete set of parameters characterizing a peak in a GMM. If the peaks overlap to a high degree and no informative SNV genotypes are available, reads are instead randomly assigned to peaks.
9. Final copy number genotype estimates, consensus sequences, and copy number confidence intervals are computed. *If the k-mer functionality is enabled, motif-sized k-mers are computed for each allele*.
10. A final report is generated with STR genotypes, parameters used, peak assignments (and peak assignment method used) for each read, copy numbers determined for each read, standard deviations for peaks, and optionally:

a. Read-level SNV calls.
b. Read- and/or peak-level motif-size k-mer quantification.

### Benchmarking genome-scale tandem repeat genotyping

We first obtained unaligned HiFi and ONT R10 Duplex LRS data for the HG002 individual (see **Availability of data and methods**). We aligned these reads to the UCSC HG38 analysis set reference genome. The reads were aligned using `minimap2` v2.28, using the `map-hifi` and `lr:hq` presets for the HiFi and ONT R10 duplex data, respectively.

English *et al*. (2024) have published a genome-wide tandem repeat benchmark via the Genome-in-a-Bottle consortium (GIAB) for the HG002 Ashkenazim individual. We used version 1.0 of this benchmark, which includes 1 784 804 regions, some with multiple tandem repeat loci defined in the same region. The authors extended the `Truvari` software (English *et al*., 2022), which can be used to benchmark structural variant calls, to support region-based STR calls. For benchmarking, we used `Truvari` version 4.3.1. We filtered out some complex or non-STR tandem repeat definitions from their benchmark region catalogue, using the following steps:

1. Motifs were normalized (e.g., CACACA → CA).
2. Homopolymers and tandem repeats with motif length >10bp were removed.
3. Benchmark regions with more than one tandem repeat were removed, since unphased proximate haplotypes cannot be correctly evaluated with `Truvari`.

Additionally, homopolymers were filtered from the HG002 benchmark variant VCF to mitigate erroneous false negatives when homopolymers were present near other STR regions. To combine and visualize the benchmarking results after running `Truvari`, we used `Laytr` (commit `c2eccbf`; https://github.com/ACEnglish/laytr). Applying this filtering, we ended up with a catalogue of 914 676 loci.

For benchmarking `STRkit`, we chose four recently-developed general-purpose LRS STR genotyping tools, all of which take standard BAM/CRAM-formatted alignment files and can output VCFs, for use with `Truvari`. The set of tools, with their versions used for benchmarking, are as follows:

- `LongTR` v1.1 (Ziaei Jam *et al*. 2024)
- `Straglr` v1.5.3 (Chiu *et al*. 2021) with `TRF` v4.09.1 (Benson 1999)
- `STRkit` v0.20.0 (*our method, benchmarked with and without SNV incorporation*)
- `STRdust` (De Coster *et al*. 2024) commit `3f3ebf0` (Aug. 22, 2024)
- `TRGT` v1.4.1 (Dolzhenko *et al*. 2024)

Parameters used for benchmarking can be found in the tool batch scripts located at https://github.com/davidlougheed/strkit_paper/tree/main/2_giab_calls. This folder also contains scripts used to generate **Figure S1** and compute the summary data found in **Table 2** and supplemental tables.

To calculate the approximate phasing error rate in `STRkit`’s SNV calling approach, we compared multi-locus phase sets derived from SNVs with haplotype-resolved output from `STRkit` (via the `--use-hp` flag) using haplotype-tagged aligned PacBio HiFi reads. A phase set was counted as containing an error if heterozygous loci did not match in terms of allele size or copy number ordering.

### Assessing computational performance

All STR genotyping tools were run on the Digital Research Alliance of Canada’s Béluga cluster, using 8 cores of an Intel Gold 6148 Skylake processor, or 1 core where parallel processing was not supported (i.e., `LongTR`). Compute time was normalized into CPU-minutes by multiplying runtime by the number of cores used.

### Calculating Mendelian inheritance in call sets

We first obtained unaligned HiFi reads for the Ashkenazi trio from PacBio. We aligned these reads to the same UCSC HG38 analysis set used for benchmarking, using `minimap2` v2.28 with the `map-hifi` preset.

In `STRkit`, we include a module named `strkit mi` to calculate rates of trio Mendelian inheritance (MI) in autosomal loci from the output of all STR genotyping tools we benchmarked. We calculated MI in terms of binary ‘yes/no’ inheritance, inheritance ±1 repeat unit, and parent-offspring 95% confidence interval overlap. The tool performs the following steps:

1. For each locus in the STR callset for the child in the trio, check if a corresponding call can be found in both parent callsets. If not, skip the locus.
2. If the locus is in a user-specified exclusion set, e.g., for removing known *de novo* mutation, skip it.
3. Add this locus to the set S of seen loci for this trio.
4. If (slightly different from step 1) a call *failed* in any of the three individuals, skip the locus (after it has been added to set S.)
5. If the calls for the locus are consistent with Mendelian inheritance (for each version of the MI metric X ∈ {strict, ±1 repeat, 95% CI, sequence, seq. len., seq. len. ±1 base pair}), add one to a counter c_X_ of consistent loci for metric X. Some versions of the metric are not supported by some of the genotyping methods; `LongTR` and `STRdust` do not provide copy numbers or 95% copy number confidence intervals, and `Straglr` does not provide 95% copy number confidence intervals or consensus allele sequences.
6. After all loci have been processed, return the total rate of MI as c/|S|.

### Genotyping expansions from targeted sequencing data

We accessed a dataset of Pacific Biosciences targeted sequencing of expansions from the following URL: https://downloads.pacbcloud.com/public/dataset/RepeatExpansionDisorders_NoAmp/. The same STR genotyping tool versions used in our benchmarking were compared with this targeted disease expansion sequencing dataset. Batch scripts with specific parameters, as well as the script used to generate **Figure 3**, are available at https://github.com/davidlougheed/strkit_paper/tree/main/4_pathogenic_exp. Coriell genotypes were manually collected from https://www.coriell.org/ Oct 4, 2022.

## DECLARATIONS

### Ethics approval and consent to participate

Not applicable.

### Consent for publication

Not applicable.

### Availability of data and materials

The source code for the STRkit package is available on GitHub at: https://github.com/davidlougheed/strkit/ and in Zenodo under the DOI https://doi.org/10.5281/zenodo.12689906, and is obtainable as a Python package in the Python Package Index (PyPI) under the name strkit.

The Genome-in-a-Bottle (GIAB) Tandem Repeats V1.0 benchmark is available from the NCBI at:

https://ftp-trace.ncbi.nlm.nih.gov/ReferenceSamples/giab/release/AshkenazimTrio/HG00 2_NA24385_son/TandemRepeats_v1.0/GRCh38/. Corresponding benchmark region annotations are available from Zenodo at https://doi.org/10.5281/zenodo.6930201.

The Genome-in-a-Bottle Ashkenazi trio sequencing HiFi data (both unaligned and aligned phased reads) used for benchmarking and Mendelian inheritance analysis are available from PacBio at: https://downloads.pacbcloud.com/public/revio/2022Q4/.

Oxford Nanopore R10 stereo duplex sequencing data for the same trio are available from the Human Pangenomics Project at: https://human-pangenomics.s3.amazonaws.com/index.html?prefix=submissions/0CB93 1D5-AE0C-4187-8BD8-B3A9C9BFDADE--UCSC_HG002_R1041_Duplex_Dorado/Dorado_v0.1.1/stereo_duplex/.

The HG38 human reference genome (analysis version) is available from UCSC at: https://hgdownload.soe.ucsc.edu/goldenPath/hg38/bigZips/analysisSet/hg38.analysisSe t.fa.gz.

Targeted expansion sequencing data used for assessing expansion genotyping are available from PacBio at: https://downloads.pacbcloud.com/public/dataset/RepeatExpansionDisorders_NoAmp/. Coriell genotypes used for assessing expansion genotyping were manually collected from https://www.coriell.org/ Oct 4, 2022.

The scripts used to analyze the data for this manuscript are available in GitHub at https://github.com/davidlougheed/strkit_paper/.

## Authors’ contributions

D.R.L. developed the software package and performed the benchmarking analyses. G.B. and D.R.L. conceived of the project. T.P. gave guidance and insight during the development of the project. G.B. supervised the project. All authors read and approved the final manuscript.

## Acknowledgements

We would like to thank Rob Eveleigh and Mathieu Bourgey for providing valuable instruction on long read data and benchmarking approaches, José Héctor Gálvez López for his important feedback on our genotyping approach, Elia Afanasiev for his help increasing computational performance and for proofreading the manuscript, and Boriana Apostolov and Stephen C. Lougheed for proofreading the manuscript. We would like to acknowledge Calcul Quebec, SecureData4Health and the Digital Research Alliance of Canada for computing resources.

