## Supplementary Material for "STRkit: precise, read-level genotyping of short tandem repeats using long reads and single-nucleotide variation"

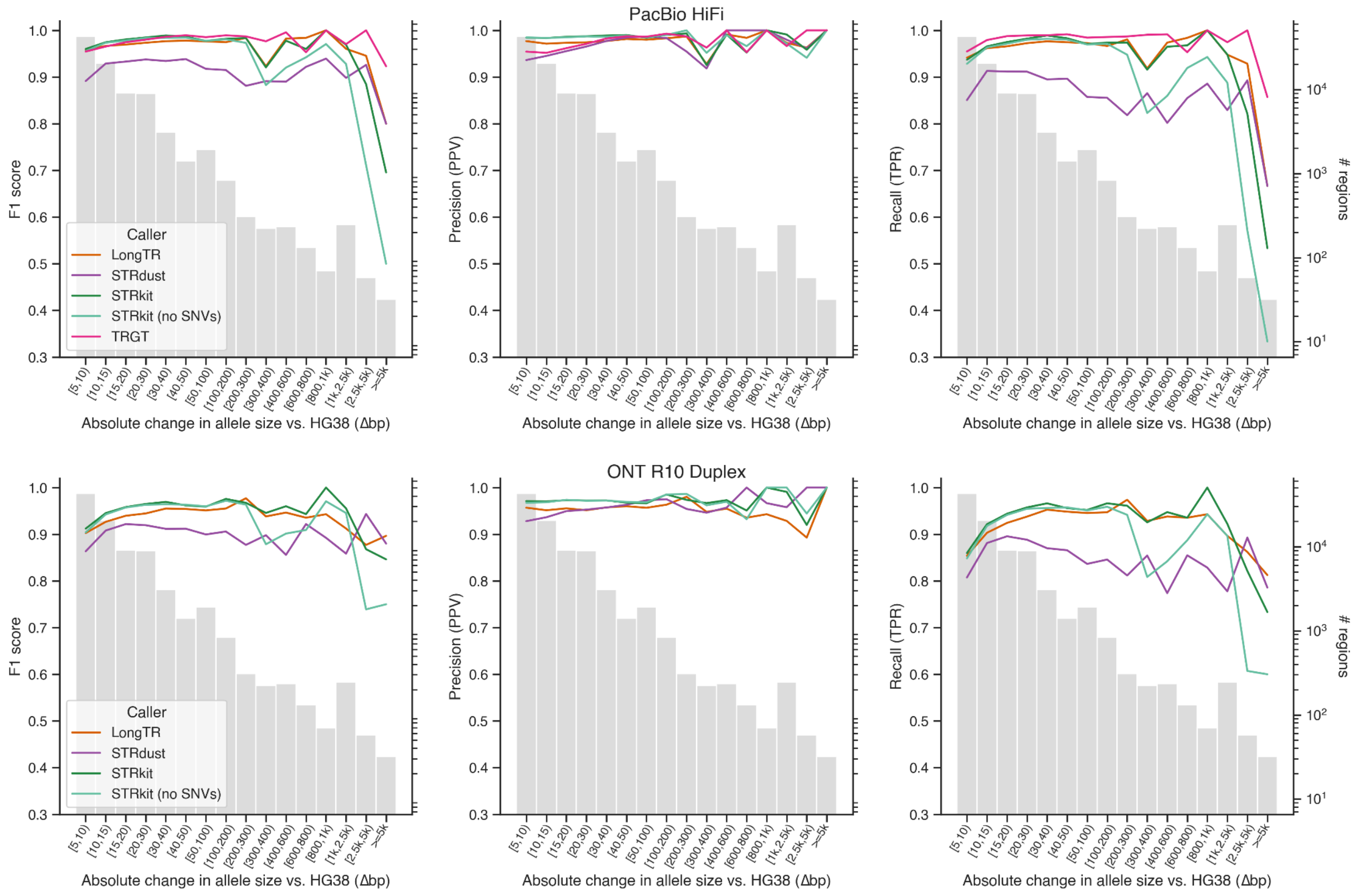


**Figure S1:** F1 score, precision, and recall of STR genotyping tools by locus-maximum allele size delta, i.e., the maximum change in allele size versus the HG38 reference genome of the maternal and paternal alleles for each locus, as output by the `Truvari` benchmarking utility and `Laytr` reporting tool.

**Table S1:** `STRkit` STR allele peak-calling method for Genome-in-a-Bottle Ashkenazi trio samples

| Tech. | Sample | SNV | SNV+Dist | Dist | Single | Not Called | % SNV use |
|---|---|---|---|---|---|---|---|
| HiFi | HG002 | 37.72% | 33.93% | 25.07% | 3.16% | 0.12% | 71.65% |
| | HG003 | 33.39% | 35.38% | 27.96% | 3.16% | 0.12% | 72.13% |
| | HG004 | 35.75% | 35.95% | 27.69% | *N/A** | 0.54%* | 71.70% |
| ONT | HG002 | 52.88% | 19.68% | 24.13% | 3.10% | 0.22% | 72.55% |

`STRkit` STR allele peak-calling method for Genome-in-a-Bottle Ashkenazi trio samples. “SNV” means 2+ SNVs were used to peak-call alleles. “SNV+Dist” means a combined one-SNV and copy number distance metric was used. “Dist” means only copy-number distance was used. “Single” means the locus was called as haploid (i.e., the X/Y-chromosomes for standard chromosomal males).

*HG004 is the maternal parent in the trio, with a standard XX karyotype and thus no haploid chromosomes (i.e., single-peak genotypes) and no Y-chromosome loci called.

**Table S2:** Rates of Mendelian inheritance (MI) for STR genotypers on the Ashkenazi trio

| Caller | MI % (copy num.) | MI % (seq.) | MI % (seq. len.) | MI% (seq. len. ±1bp) | # trio calls |
|---|---|---|---|---|---|
| **`STRkit`** | 98.53% | 97.85% | 98.07% | 99.08% | 884 009 |
| **`STRkit (-SNV)`** | 98.67% | 93.65% | 95.68% | 98.43% | 883 888 |
| **`LongTR`** | *N/A* | 94.52% | 94.85% | 97.54% | 882 515 |
| **`Straglr`** | 86.14% | *N/A* | *N/A* | *N/A* | 854 887 |
| **`STRdust`** | *N/A* | 69.56% | 71.55% | 92.88% | 882 322 |
| **`TRGT`** | 98.29% | 94.68% | 97.10% | 98.98% | 883 449 |

Rates of Mendelian inheritance (MI) for STR genotypers on the Ashkenazi trio with HiFi sequencing data, using regions from the Genome-in-a-Bottle tandem repeats v1.0 benchmark, as measured by `STRkit`'s Mendelian inheritance calculator. Up to four different versions of the MI metric are measured, depending on the caller: copy number MI (exact MI in terms of [approximate or reported] copy number), sequence MI (exact MI in terms of allele sequence), sequence length MI (exact MI in terms of allele length, in base pairs), and sequence length MI ±1 base pair. The "# trio calls" figure indicates the number of loci with successful calls in all three individuals of the trio, out of a total of 914 676. Underlines indicate the best value for the Mendelian inheritance metric, within the sequencing technology.

**Table S3:** Runtime performance and maximum memory usage of STR genotyping software

Panel A: Runtime (core-minutes)

| | PacBio HiFi (Coverage: 32.3x, 31.6x, 32.6x) | | | | ONT R10 Duplex (~12x) |
|---|---|---|---|---|---|
| **Caller** | **HG002** | **HG003** | **HG004** | **Avg. Runtime** | **HG002** |
| `STRkit` | 494 | 492 | 977 | 654 | 492 *(best multi-core)* |
| `STRkit (-SNV)` | 977 | 976 | 980 | 978 | 492 (") |
| `LongTR` | 244 | 184 | 244 | 224 *(best)* | 184 *(best)* |
| `Straglr` | 4396 | 3907 | 4395 | 4233 | 3911 |
| `STRdust` | 11231 | 9765 | 11227 | 10741 | 9277 |
| `TRGT` | 373 | 356 | 406 | 378 *(best multi-core)* | *N/A* |

Panel B: Maximum memory usage (gigabytes [GB])

| Caller | HG002 | HG003 | HG004 | Avg. Max. Memory | HG002 |
|---|---|---|---|---|---|
| `STRkit` | 10.7 GB | 11.1 GB | 10.1 GB | 10.6 GB | 7.3 GB |
| `STRkit (-SNV)` | 6.5 GB | 6.4 GB | 6.6 GB | 6.5 GB | 4.3 GB |
| `LongTR` | 3.1 GB | 2.7 GB | 3.1 GB | 3.0 GB | 5.5 GB |
| `Straglr` | 78.6 GB | 77.1 GB | 78.9 GB | 78.2 GB | 35.9 GB |
| `STRdust` | 12.9 GB | 13.1 GB | 10.4 GB | 12.1 GB | 14.7 GB |
| `TRGT` | 1.3 GB | 2.8 GB | 1.1 GB | 1.7 GB | *N/A* |

Runtime performance and maximum memory usage of STR genotyping software on our GIAB tandem repeats benchmark subset for the Ashkenazi trio. Underlines indicate the best value for the performance metric, within the sequencing technology. *N/A* is not available because of licensing restrictions.

### **Table S4:** HTT and FMR1 expansion genotypes from long-read STR calling tools

Panel A: HTT genotyping results

| Sample | Documented genotype* | `STRkit` | `LongTR`[1] | `Straglr` | `STRdust`[1] | `TRGT`[1] |
|---|---|---|---|---|---|---|
| NA13505 | 22/**50** | 22/**51** | 22/**51** | 22/**51** | 23/**52** | 23/**53** |
| NA13509 | 15/**70** | 16/**75** | 15/**74** | 15/**75** | 15/**75** | 20/**76** |
| NA20253 | 22/**96-103**[a] | 22/**114** | 22/**113** | 22/**111** | 23/**121** | 23/**111** |
| NA14044 | 19/**250**[b] | 19/**804** | 19/**723** | 20/*20* | 20/**1210** | 22/**646** |
| HEK293 | Control (no expansion) | 17/18 | 17/18 | 17/17 | 17/18 | 18/19 |

* The `Tandem Repeats Finder` (`TRF`) catalogue includes a tailing CAACAG as part of the HTT repeat, so we subtracted 2 from reported repeat counts for comparison against the documented genotypes.

[1] These tools occasionally included additional non-CAA/CAG repeats in the VCF output for the locus. We identified the CAG stretch in the reported expanded allele and calculated the copy number.

[a] Mean PCR genotype across 10 volunteer laboratories from Kalman *et al.* 2007. Multiple peaks are visible in this sample, which De Luca *et al.* (2021) also found via repeat-primed PCR.

[b] In this sample, there is a wide range of copy numbers in the expanded allele (i.e., mosaicism or somatic instability) visible in the read-level data.

Panel B: FMR1 genotyping results

| Sample | Documented genotype† | `STRkit` | `LongTR`[1] | `Straglr` | `STRdust`[1] | `TRGT` |
|---|---|---|---|---|---|---|
| NA13664 | 28±3/49±3 | 31/**54** | 31/**53** | 32/**54** | 32/**53** | 32/**54** |
| NA06896 | 23/**95-140**[c] | 24/**187** | 24/**196** | 24/**196** | 24/**150** | 24/**192** |
| NA07537 | 28-29/**>200** | 30/**343** | 29/**348** | 30/**341** | 29/*29* | 30/**339** |
| HEK293 | Control (no expansion) | 30/31 | 30/32 | 32/32 | 30/30 | 31/32 |

† The `TRF` catalogue includes an interrupting section (4 amino acids) in the FMR1 repeat which we subtracted from reported repeat counts in order to match the Coriell genotype.

[c] Pre-mutation expansion range (linked to FXPOI and FXTAS).

HTT and FMR1 genotypes from long-read STR calling tools using high-depth targeted CCS in seven samples with expansions plus a control sample. Pathogenic-length expansions are shown in bold text; expansions at the upper limit of "normal" are underlined; expansion alleles which were missed are italicized. Documented sample genotypes are from the Coriell institute, except where noted; accessed from https://www.coriell.org/ Oct 4, 2022.
